# Contingency violation in extinction learning

**DOI:** 10.1101/2025.04.28.651099

**Authors:** Robert Willma, Juan Peschken, Roland Pusch, Jonas Rose

## Abstract

In extinction learning, contextual renewal occurs when an extinguished behavior reemerges after a context change. A key question is how stimuli become integrated as contextual cues. While contingency, the predictive relationship between stimuli and outcomes; is known to be important, its precise contribution remains unclear. Using a ABA renewal design with pigeons in operant chambers, we systematically violated contingency by probabilistically reinforcing responses during extinction. Our results show that partial violations of contingency modulated extinction learning but did not abolish contextual renewal. Instead, pigeons developed meta-learning strategies, adapting their behavior across sessions to optimize reward despite extinction conditions. These findings highlight that context formation is sensitive to contingency levels, but also that animals can flexibly reorganize their learning strategies when contingency is unstable.

## Introduction

Life unfolds in an environment saturated with stimuli, where organisms must constantly filter information to prioritize what is relevant for survival. However, survival depends not only on prioritizing relevant stimuli but also on flexibly updating these selections when environmental conditions change. Organisms must recognize when previously relevant information becomes obsolete and adjust their behavior accordingly. Extinction learning is central to this process, diminishing previously established associations when expected outcomes no longer occur. Traditionally, extinction has been understood as the inhibition of learned responses rather than their erasure, as evidenced by the renewal effect, where extinguished behaviors reappear when an individual encounters a different context after extinction.

A classical view in learning theory distinguishes between stimuli that directly guides behavior, known as cues, and those that provide the background conditions for learning, referred to as context. For instance, a pigeon might reliably find seeds beneath a specific tree, making the tree a salient cue that predicts reward availability. However, broader environmental features (i.e. the season, light conditions, and temperature) constitute the context within which this cue operates. When winter arrives, the tree itself remains unchanged, but shifts in the surrounding context signal that food is no longer available. This example highlights how behavioral control depends not only on the direct association between a cue and an outcome, but also critically on the contextual framework in which this association is embedded.

While many studies have portrayed context, as in the previous example, as a passive collection of environmental features surrounding learning events, recent research challenges this view. Rather than being a static backdrop, context appears to emerge dynamically through competition among stimuli present during learning. This competitive framework suggests that stimuli with higher associative relevance, determined by factors such as contiguity and salience, are more likely to serve as the context controlling behavior. However, this is not the complete picture. A critical factor in any form of associative learning is contingency.

Contingency, defined as the reliability with which a stimulus predicts outcomes, plays a central role because it defines the predictive value of contextual information. A stimulus that consistently co-occurs with an outcome provides more informative value and is therefore more likely to be integrated as part of the context. In the pigeon example, the seasonal context has a high contingency with the availability of food. If random food deliveries began occurring during winter, the predictive value of the winter context would diminish. If context is fundamentally constructed through learning mechanisms, then the capacity to flexibly update behavior during extinction should depend heavily on how organisms evaluate the predictive relationships between stimuli and outcomes.

Building on earlier work that established the importance of competition among stimuli— particularly the roles of contiguity and salience during extinction and renewal—the present study focuses specifically on the role of contingency. We employed a previously reported task using Skinner boxes, following a classical ABA paradigm but involving only a single context. By isolating contingency as a variable within one context, we aim to determine how violations of contingency affect context representations and, in turn, control the renewal of conditioned behaviors. Our objective is to determine the level of contingency necessary for a stimulus to be integrated and function as a context, and, conversely, the threshold at which a stimulus is disregarded and fails to act either as a context or as a cue.

## Materials and methods

### Subjects

Six experimentally naïve pigeons (*Columba livia*) of unknown age were used in this study. The birds were sourced from local breeders and housed individually in a colony room. Throughout the experiment, water was available ad libitum, and a controlled feeding schedule was implemented. Birds earned their food rewards during the sessions, with additional feeding provided if needed afterward. All experimental procedures complied with German animal welfare regulations, adhered to the European Communities Council Directive 86/609/EEC, and were approved by the ethics committee of the State of North Rhine-Westphalia, Germany.

### Experimental setup

[To be completed]

### Behavioral protocol

[To be completed]

### Statistical analysis

[To be completed]

## Results

[Draft text – will be finalized]

We assessed the behavior of six pigeons engaged in a two-stimulus discrimination task in operant chambers, using a within-session acquisition-extinction-renewal (ABA) design. To examine how contingency affects contextual control over extinguished behavior, we systematically violated the reliability of the extinction context by delivering food rewards probabilistically at varying reinforcement ratios.

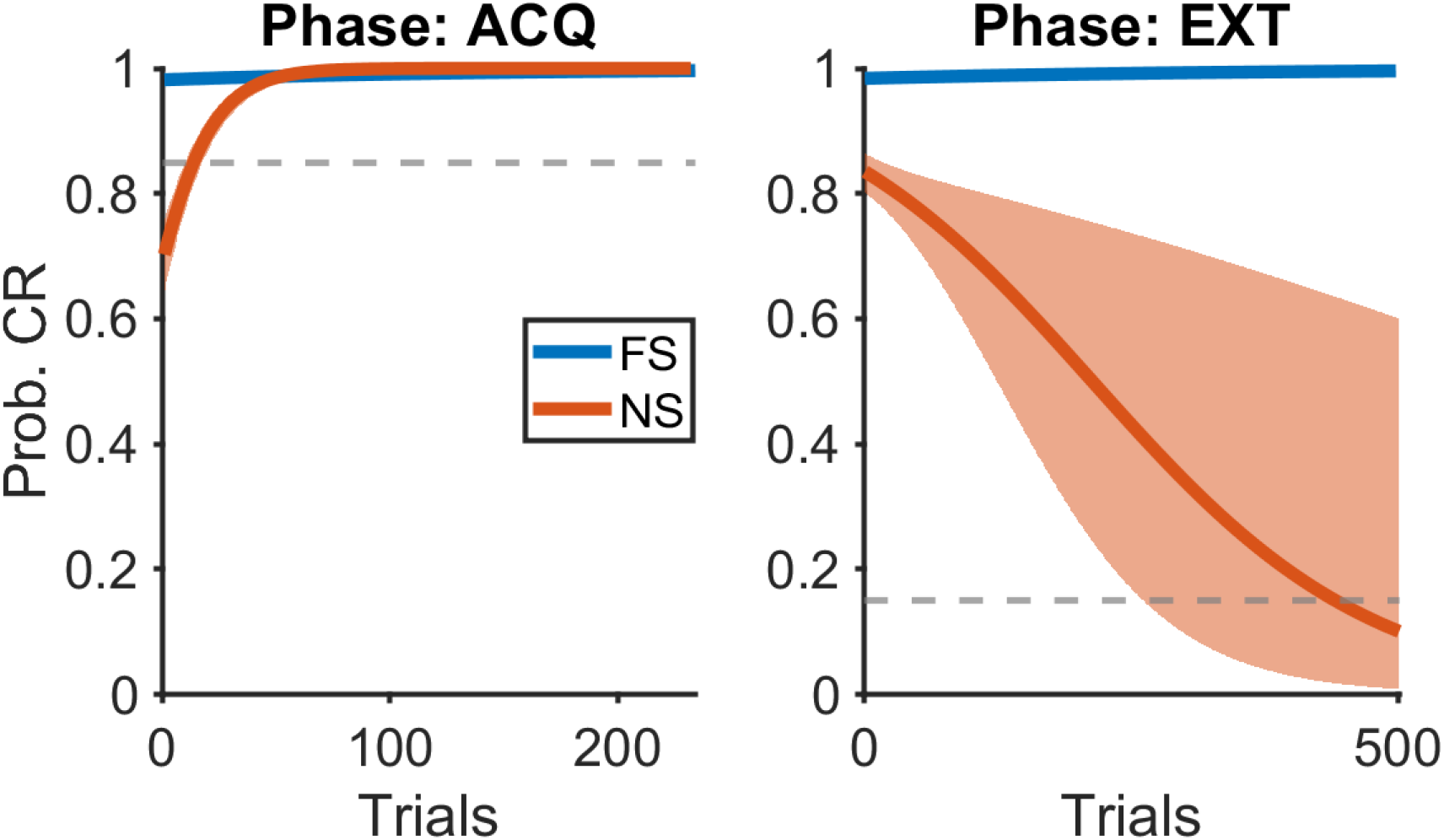
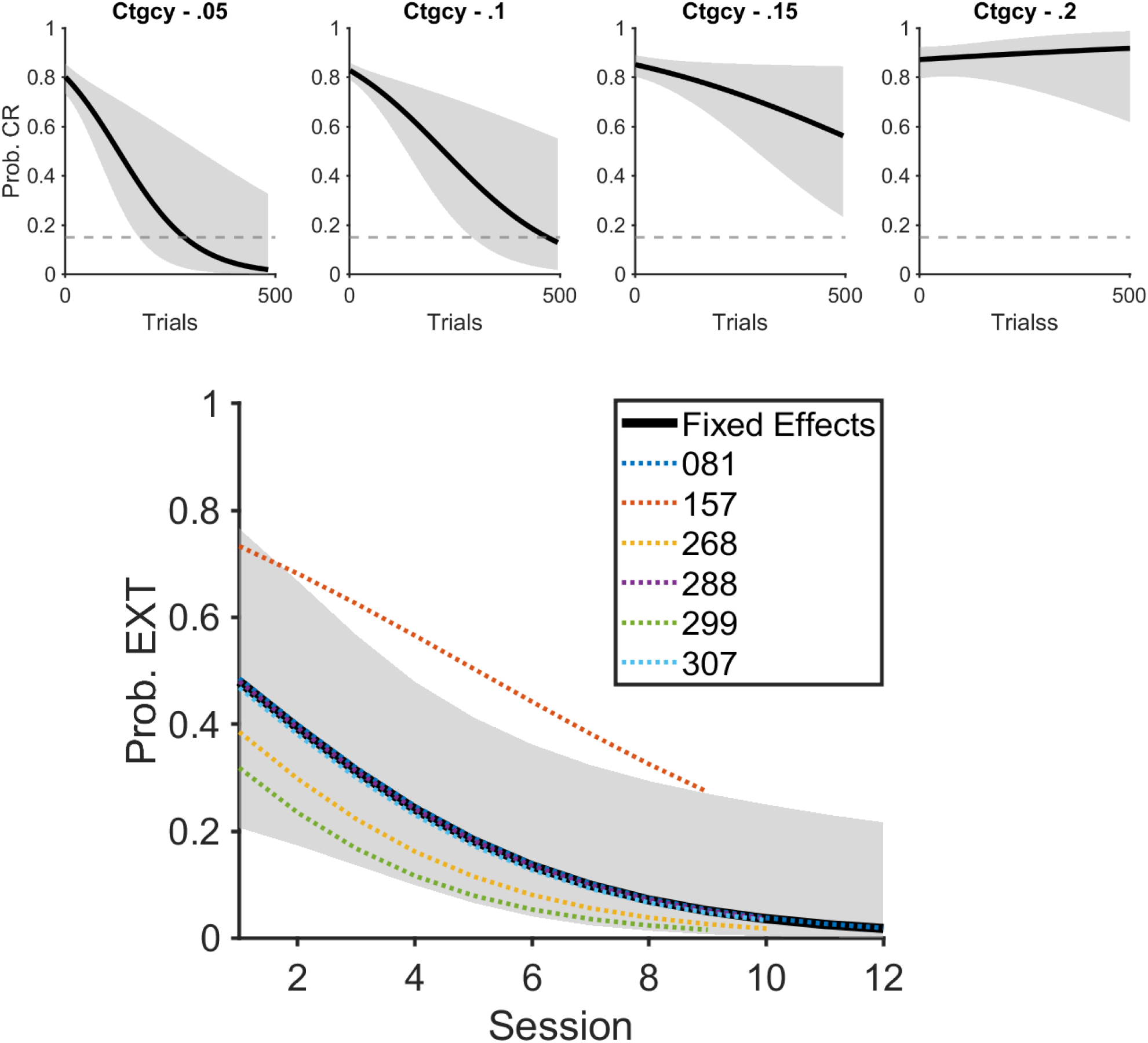

## Discussion

[Draft text – will be finalized]

A simple explanation for the continued pecking during the extinction phase might be that trial length was not controlled. A reaction to a novel stimulus could have advanced the session faster, leading to a familiar, rewarded stimulus appearing sooner. To maximize food intake, pigeons had to remain in the acquisition phase by maintaining performance below an 85% success rate, although this threshold was unknown to them. Consequently, pigeons continued to peck even after entering the extinction phase, where novel stimuli were rarely reinforced but familiar stimuli still appeared. However, similar designs with intermixed reinforced and unreinforced trials (Packheiser, Güntürkün & Pusch, 2019; Peschken, 2025) did not show the same persistent behavior. The key difference here was the intermittent reinforcement of novel stimuli, a factor that future studies could control more systematically.

### Intermittent Reinforcement as a Protective Factor

Intermittent reinforcement has long been known to maintain behavior even under low reinforcement rates (Ferster, 1960; Eckerman & Lanson, 1969). In fact, such schedules often produce more resilient behavior than continuous reinforcement (Zarcone et al., 2013; Roberts, Bullock & Bitterman, 1963; Kendall, 1974). Here, the lowest reinforcement rate used (5%) was expected to promote extinction, whereas the highest (20%) was designed to maintain responding. Prior pigeon studies have employed rates as low as 12.5% without achieving extinction (Roper & Zentall, 1999; Roberts, MacDonald & Lo, 2018; Stagner & Zentall, 2010; Zentall & Laude, 2013). Similarly, Stahlman, Young, and Blaisdell (2010) showed that pigeons continue to peck even when reinforcement probabilities drop as low as 0.6%, though lower probabilities produced fewer responses. Notably, in both their work and ours, low- and high-reinforcement trials were presented in the same session. The presence of reliably rewarded trials could protect against extinction by maintaining general motivation. Peschken et al. (2025) demonstrated extinction in a similar design, but using a large arena, where pigeons could avoid an extinguished stimulus physically—unlike the forced exposure within the confines of a Skinner box.

### Evidence for Meta-learning and Retroactive Metacognition

The pigeons’ adaptation across sessions suggests a form of meta-learning. Computational models describe learning within two loops: a fast, within-session loop and a slower, across-sessions loop (Nussenbaum & Hartley, 2024). In our task, pigeons initially extinguished novel stimuli within sessions, but with experience, extinction became less likely. This shift parallels meta-cognitive control, where subjects monitor and adjust their behavior based on accumulated experience (Nelson, 1990). Although animal metacognition remains debated (Smith, 2009; Carruthers, 2008), the pigeons’ behavior—persistently pecking to prolong access to rewarding trials—suggests strategic use of reward history. Similar abilities have been shown in pigeons during working memory tasks requiring prospective metacognitive judgments (Castro & Wasserman, 2012; Iwasaki, Watanabe & Fujita, 2017) and retroactive metacognitive processes (Shimp, 1982; Adams & Santi, 2010).

### Responses to Novel Stimuli

Even though extinction was not achieved in most sessions, pigeons’ responses to novel stimuli differed from familiar ones. If novel stimuli had been treated as completely unrewarding (like familiar negatives), choice behavior should have been random, yet this was not observed. Reaction times to peck novel stimuli were slower, particularly during extinction phases, suggesting hesitation or evaluation processes before responding. This delay could reflect cognitive uncertainty. In addition, peck dispersion was higher for novel stimuli, consistent with reports that lower reinforcement rates lead to less focused pecking behavior (Stahlman, Young & Blaisdell, 2010). Familiar stimuli, extensively trained, elicited more precise pecks, reflecting pigeons’ strong visual learning capabilities (Vaughan & Greene, 1984; Wasserman, Turner & Güntürkün, 2024; Pusch, Clark, Rose, & Güntürkün, 2023).

## Conclusion

Manipulating contingency through rare intermittent reinforcement prevented extinction of pecking behavior. Delivering rewards in just 5% of “extinction” trials was sufficient to sustain responding. Over time, pigeons exhibited meta-learning by adapting their behavior to maximize rewards across sessions. Rather than “successfully” extinguishing, which ended the session early, pigeons maintained behavior, thereby optimizing their food intake under the given task dynamics.

